# Engineered *Bacillus subtilis* as oral probiotics to enhance clearance of blood lactate

**DOI:** 10.1101/2023.11.30.569300

**Authors:** Mengdi Yang, Noah Hutchinson, Ningyuan Ye, Jianing Yin, Ming Guan, Zongqi Wang, Peiru Chen, Shaobo Yang, Justin D. Crane, Ke Zhang, Xuesong He, Jiahe Li

## Abstract

Elevated lactate concentrations are implicated in various acute and chronic diseases such as sepsis and mitochondrial dysfunction, respectively. Conversely, ineffective lactate clearance is associated with poor clinical prognoses and high mortality in these diseases. While several groups have proposed using small molecule inhibitors and enzyme replacement to reduce circulating lactate, there are few practical and effective ways to manage this condition. Recent evidence suggests that lactate is exchanged between systemic circulation and the gut, allowing bidirectional modulation between the gut microbiota and peripheral tissues. Inspired by these findings, this work seeks to engineer spore-forming probiotic *B. subtilis* strains to enable intestinal delivery of lactate oxidase as a therapeutic enzyme. After strain optimization, we showed that oral administration of engineered *B. subtilis* spores to the gut of mice reduced elevations in blood lactate in two different mouse models involving exogenous challenge or pharmacologic perturbation without disrupting gut microbiota composition, liver function, or immune homeostasis. Taken together, through the oral delivery of engineered probiotic spores to the gastrointestinal tract, our proof-of-concept study offers a practical strategy to aid in the management of disease states with elevated blood lactate and provides a new approach to ‘knocking down’ circulating metabolites to help understand their roles in host physiological and pathological processes.

**Significance Statement:** This study pioneers the use of engineered *Bacillus subtilis* spores as an oral probiotic therapy to enhance the clearance of elevated blood lactate, a condition linked to severe health issues like sepsis and metabolic disorders. By genetically modifying these spores to deliver therapeutic enzymes directly to the gut, we demonstrated a practical, effective method to modulate systemic lactate levels. This approach leverages the natural exchange between the gut microbiota and systemic circulation, offering a new strategy for managing diseases associated with lactate dysregulation. The safety and efficacy of this method were validated in mouse models, providing a foundation for future clinical applications.

## Introduction

Lactate is a hydroxycarboxylic acid and a byproduct of anaerobic glycolysis. When cellular energy demands exceed the rate of oxidative phosphorylation, glycolysis is upregulated to rapidly produce adenosine triphosphate (ATP), and cytosolic pyruvate is converted to lactate by lactate dehydrogenase (LDH) (1). In addition to its role in ATP production, the LDH reaction plays an essential role in the regulation of pH and cellular redox through the oxidization of NADH and the assimilation of protons. Since this reaction is reversible and susceptible to concentration-dependent regulation by its products, the ubiquitous expression of monocarboxylate transporters (MCTs) enables extracellular and systemic lactate concentrations to influence cellular metabolism, acidity and NAD^+^/NADH ratios (2) (3). As such, maintaining lactate homeostasis is crucial for the regulation of various biological functions (1). While lactate influxes serve as crucial regulatory signals in many physiologic processes, uncontrolled accumulation of lactate, coined lactic acidosis, can cause physical immobilization, organ failure, hypoxia, and septic shock (4). Further, chronic exposure to elevated lactate concentrations can cause metabolic abnormalities, immune dysregulation (5), hindered wound healing (6), and cellular senescence (2), ultimately contributing to the development of cancer, physical frailty, cardiovascular disease, diabetes, and neurodegeneration (7, 8). Given these implications, strategies to accelerate the clearance of excess lactate and promote the maintenance of healthy blood lactate values possess great utility in the clinical setting. Various lactate-centric treatment methods have been developed, including oxygen therapy (9), intravenous fluid replenishment (10), and medications targeting LDH (11) or lactate transporters (12). However, most of these methods do not directly target lactate itself.

Recently, a new strategy was developed that directly mitigates lactic acidosis and intracellular reductive stress by intravenously delivering a dual-enzyme system composed of lactate oxidase (LOX) and *Escherichia coli* catalase (CAT), which irreversibly converts extracellular lactate and oxygen to pyruvate and water (13) (**Figure 1A**). While effective, the direct injection of bacterial enzymes into the bloodstream faces several challenges for translation associated with potential immunogenicity, short serum half-life, and practical considerations such as cost, storage conditions, and difficulties surrounding their requirement for frequent dosing (13). Recent evidence indicates that lactate can freely exchange between peripheral tissues, circulation, and the gut lumen, promoting bidirectional modulation between one another (14–16) (**Figure 1B**). This relationship, combined with bacterial engineering techniques and existing synthetic biology tools can be leveraged through engineered probiotics, which deliver recombinant therapeutic enzymes to the gut for the modulation of systemic lactate. To this end, we have chosen *Bacillus subtilis* as a probiotic candidate for genetic engineering because it is generally recognized as safe (GRAS) by the FDA. Furthermore, *B. subtilis* can form spores, which are associated with improved thermostability, shelf life, and survival in harsh upper gastric conditions to enable germination in the intestine (17).

**Figure 1.**
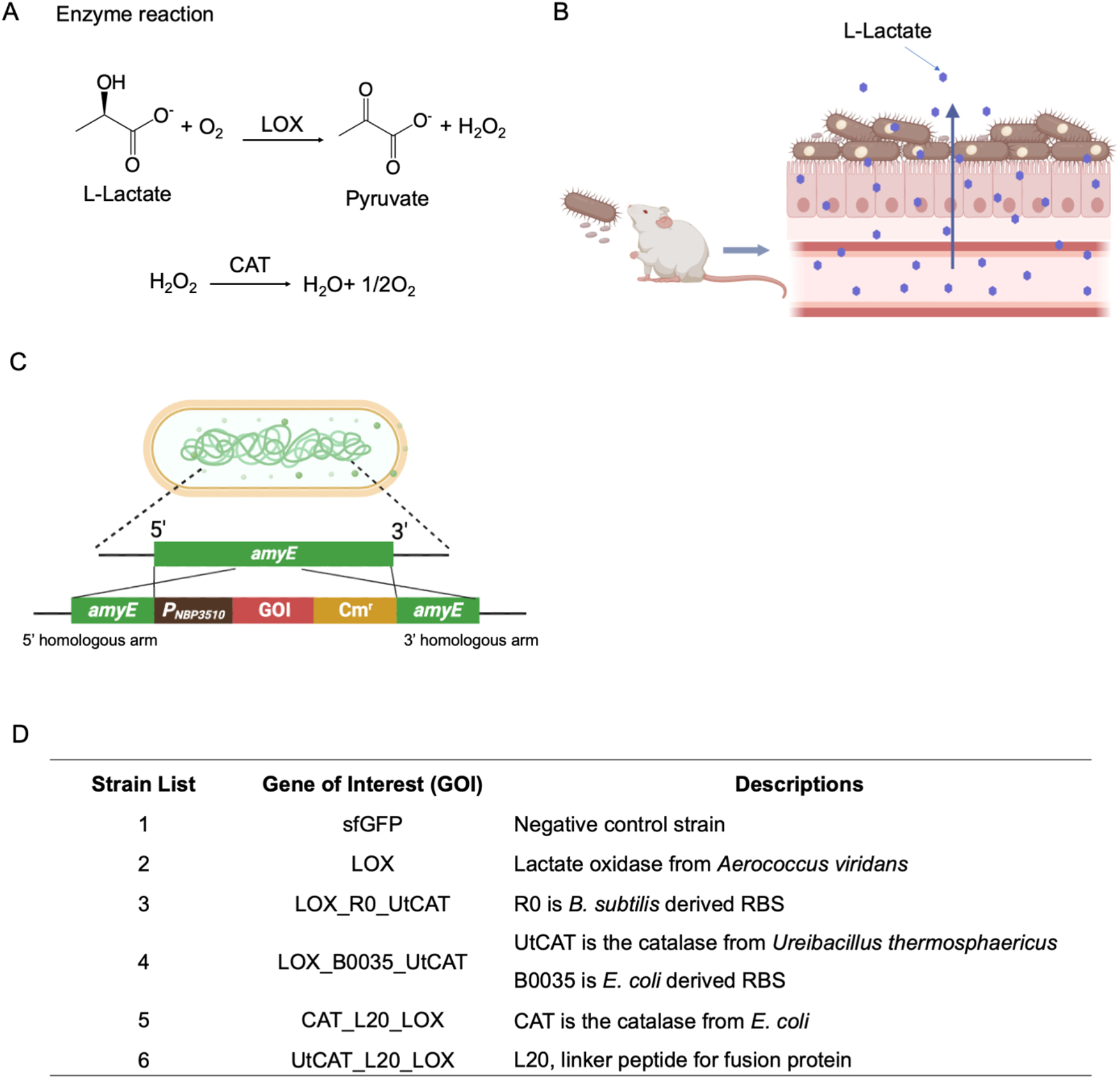
Schematic diagram and genomic integration of lactate oxidase and catalase in multiple *B. subtilis* strains. (A) Enzyme reactions of lactate oxidase (LOX) and catalase (CAT). (B) *B. subtilis* strains were engineered to express lactate oxidase (LOX), facilitating the conversion of lactate to pyruvate. (C) The genetic design involved the incorporation of constitutively expressed transgenes in a nonessential genomic locus such as the *amyE* gene in *B. subtilis*. The gene of interest (GOI) is driven by a constitutive promoter named *PNBP3510*, followed by a chloramphenicol-resistant gene (Cm^r^) for selection. The 5’ and 3’ *amyE* sequences (∼1kb in length for each side) are used for homologous recombination. (D) The list of strains used in this study encodes the following enzymes: LOX (lactate oxidase from *Aerococcus viridans*), CAT (catalase from *E. coli*), and UtCAT (catalase from *Ureibacillus thermosphaericus*). R0 and B0035 are synthetic ribosome binding sequences (RBS). L20, a 20-amino acid peptide linker between two proteins to generate fusion proteins. This figure is created using BioRender.

After genomic integration of genes encoding LOX and CAT of various configurations into different *B. subtilis* strains, we performed *in vitro* measurements of peroxide breakdown and lactate consumption to determine the optimal design. Intracellular expression of LOX in *B. subtilis PY79* was capable of efficiently clearing extracellular lactate and thus was chosen for the remainder of the study. Further mouse experiments demonstrated that oral administration of this strain in spore form was effective in not only reducing baseline blood lactate values but also mitigating lactic acidosis following exogenous administration of sodium lactate or inhibition of mitochondrial respiratory chain complex I by metformin. Additional follow-up studies indicated that the introduction of this strain in the mouse gut was safe, transient, and not disruptive to the existing microbial community. These results suggest lactate as a target molecule for efficacious removal with therapeutic potential. More broadly, our work offers an opportunity for probiotics-mediated delivery of therapeutic enzymes in the gut for the modulation of the circulating metabolome.

## Results

### Genomic integration of genes encoding lactate oxidase and catalase in multiple *B. subtilis* strains

*B. subtilis* is a Gram-positive, rod-shaped, endospore-forming bacterium isolated from soil, widely utilized for heterologous protein production (18). We initially attempted to produce recombinant enzymes extracellularly by employing the Sec-dependent secretion pathway via fusion with a signal peptide (*amyQ*) in *B. subtilis* WB800N. While WB800N is optimized for extracellular protein production with eight proteases being knocked out, it failed to produce detectable LOX in the supernatant (data not shown). It is possible that LOX (∼ 41 kDa as a monomer) can undergo tetramerization to form an even larger protein complex that prohibits efficient protein export, which is one limitation of the Sec-dependent protein secretion pathway (19). Subsequently, our focus shifted to establishing an intracellular expression system by integrating the target gene into a nonessential locus (*amyE*) in the chromosome through homologous recombination (20). To demonstrate the versatility of our approach, we sought to build and test our constructs in four different *B. subtilis* strains including PY79, WB800N, NCIB3610, and HU58 that have been employed for various preclinical trials (20–25). While PY79 and WB800N are amenable to direct transformation of DNA constructs, NCIB3610 and HU58 require phage transduction for genomic integration to allow for stable transgene expression. However, we primarily focused on PY79 in this work due to its widely available genetic information, and proven track record for safety, efficacy, and tractability (24, 26–30).

To enable high-level expression of transgenes from a single copy in the genome, we chose a highly active promoter termed *P_NBP3510_* (31). All integration fragments targeting the *amyE* locus consist of three essential components: the *amyE* 5’ homologous arm fused with *P_NBP3510_*, the gene of interest (GOI), and the *amyE* 3’ homologous arm coupled with a chloramphenicol resistance selection marker (**Figure 1C**). In pursuit of optimal catalase activity, which is necessary to break down cytotoxic hydrogen peroxide derived from LOX activity, we selected a homologous catalase from *Ureibacillus thermosphaericus*, UtCAT, which is more active and stable than the *E. coli* CAT (32). Various combinations were then created using LOX, UtCAT, and CAT (**Figure 1D**). To couple the expression of UtCAT or CAT with LOX in a single gene cassette, we explored two different configurations. The first strategy is to construct a synthetic operon consisting of the *PNBP3510* promoter with two separate ribosome binding sequences to drive the expression of LOX and UtCAT or CAT, respectively. Two strong RBSs (R0 and B0035), tailored for *B. subtilis* and *E. coli*, were found to be equally active in *B. subtilis* strains. As a result, we generated two strains: *ΔamyE::cat*_LOX_B0035_UtCAT and *ΔamyE::cat*_LOX_R0_UtCAT. In addition to the operon configuration, we also generated fusion proteins, where we inserted a linker peptide (L20) between CAT (UtCAT) and LOX, which was previously designed for intravenous administration of the fusion protein LOXCAT (13). The expression of all constructs was confirmed through Western Blot analysis (**Figure. S1**). Finally, we constructed the integration fragment incorporating the super-folding green fluorescent protein (sfGFP) in various *B. subtilis* strains as negative controls for *in vitro* and *in vivo* lactate clearance measurements (16).

### Engineered *B. subtilis* exhibited catalase and lactate oxidase activities as whole-cell biocatalysts

The activity of catalase in the CAT and UtCAT strains can be ascertained by monitoring the amount of oxygen gas generated after the direct incubation of engineered strains with hydrogen peroxide as described by others (33). Interestingly, we observed that *B. subtilis* PY79 encoding LOX or sfGFP alone exhibited the ability to break down H_2_O_2_, and the efficiencies of H_2_O_2_ clearance were comparable to those strains that were engineered to overexpress an additional copy of catalase transgene (the CAT or UtCAT) (**Figure 2A**). Since *B. subtilis* vegetative cells are known to constitutively produce catalases, which confers resistance to oxidative stress(^34^), we speculate that the native catalase activities of *B. subtilis* PY79 were sufficient to degrade H_2_O_2_, a cytotoxic intermediate from LOX, without the need to overexpress a recombinant catalase. However, it is noteworthy that the endogenous and recombinant catalase activities varied among different *B. subtilis* strains. For instance, no significant improvement of catalase activities was observed in NCIB3610 and HU58 strains that were engineered to overexpress UtCAT or CAT through either a synthetic operon or a genetic fusion configuration, in comparison to those expressing only LOX or sfGFP (**Figure S2**). Conversely, in WB800N, recombinant CAT or UTCAT enhanced catalase activities compared to LOX or sfGFP alone (**Figure S2**). Based on the above findings, we reasoned that expressing a single transgene LOX by itself would be sufficient to safely metabolize lactate without an additional copy of catalase in *B. subtilis*. As homeostatic blood lactate levels are typically < 2 mM, with levels reaching > 4 mM during severe lactic acidosis, we next chose to assess the ability of engineered strains to catabolize lactate *in vitro* with a starting concentration of lactate at 5 mM (35). Notably, PY79 expressing LOX completely removed 5 mM L-lactate within 1 hour (**Figure 2B**). To demonstrate the versatility of LOX expression in promoting the degradation of L-lactate, we further engineered three additional *B. subtilis* strains (WB800N, NCIB3610, and HU58) to express LOX, which led to complete breakdown of 5 mM L-lactate within 1-3 hours (**Figure 2C** and **Figure S3**). Based on these findings, we concluded that *B. subtilis* strains expressing intracellular LOX can directly serve as a whole-cell biocatalyst to break down extracellular lactate without the need to use purified recombinant enzymes.

**Figure 2.**
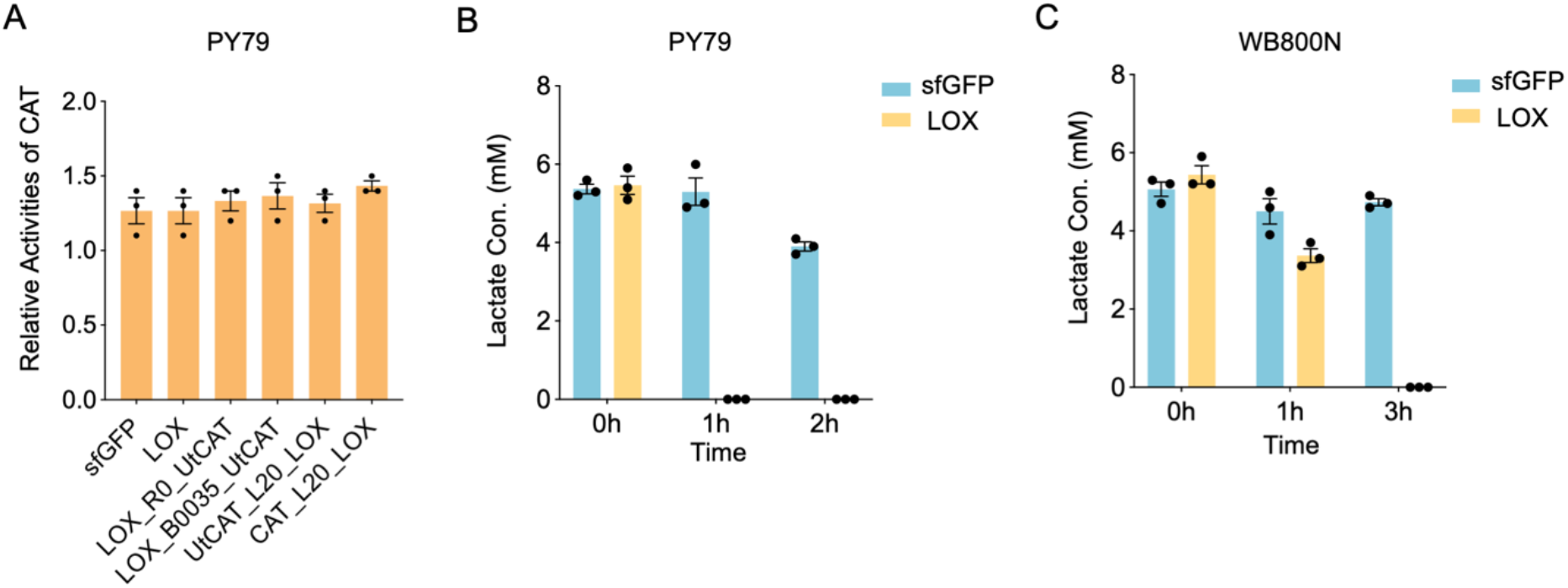
*B. subtilis* expressing LOX (*B. subtilis*^LOX^) exhibited catalase and lactate oxidase activities to clear hydrogen peroxide and L-lactate, respectively. (A) Catalase activities of *B. subtilis* PY79 strains encoding different constructs as shown in Figure 1D. *In vitro* lactate consumption of *B. subtilis*^LOX^ in PY79 (B) and WB800N (C) strains from a starting lactate concentration at 5 mM. N=3 independent experiments and n=3 technical replicates.

### Oral supplementation of engineered *B. subtilis* spores reduced baseline blood lactate and accelerated lactate clearance during metabolic challenge

Oral administration of PY79 spores has previously been shown to be safe(^24^), and the spores’ superior resistance to extreme conditions could help overcome harsh gastric conditions that diminish the viability of engineered probiotics (36). For these reasons, we further evaluated whether orally administering *B. subtilis* expressing LOX could help mitigate lactic acidosis *in vivo* (**Figure 3A**). Utilizing a published lactate induction model in mice, 6-8 week-old wild type C57BL/6 mice were randomly assigned to two groups: *B. subtilis*^LOX^ and *B. subtilis*^sfGFP^, with *B. subtilis*^sfGFP^ control serving as a bacterial load mimic which does not express LOX. Since a previous safety study in rats did not show clinically significant side effects after a 90-day daily oral administration of 10^11^ CFU/kg body weight engineered PY79 spores (24), we opted to feed the mice with the same dose (∼2x10^9^ CFU per mouse) daily for three days. On Day 3, 20 minutes after the last oral administration of spores, a baseline serum lactate level was measured before we induced experimental lactate elevation. Interestingly, it was found that *B. subtilis*^LOX^ mice displayed lower blood lactate values than *B. subtilis*^sfGFP^ (*p*=0.02, **Figure 3B**), suggesting that *B. subtilis*^LOX^ could slightly decrease homeostatic blood lactate concentrations. Subsequently, intraperitoneal injection of 2.3 g/kg body weight Na-L-lactate was administered to induce an experimental lactate spike. 30 minutes after injection, *B. subtilis*^LOX^ exhibited less potent increases in blood lactate than that of the sfGFP control group, an effect which persisted through the 75-minute timepoint post-injection before the blood lactate returned to baseline (**Figure 3C**). Of note, *B. subtilis*^LOX^ displayed a significantly reduced area under the curve (AUC, *p*=0.0027) compared to that of *B. subtilis*^sfGFP^ (**Figure 3D**).

**Figure 3.**
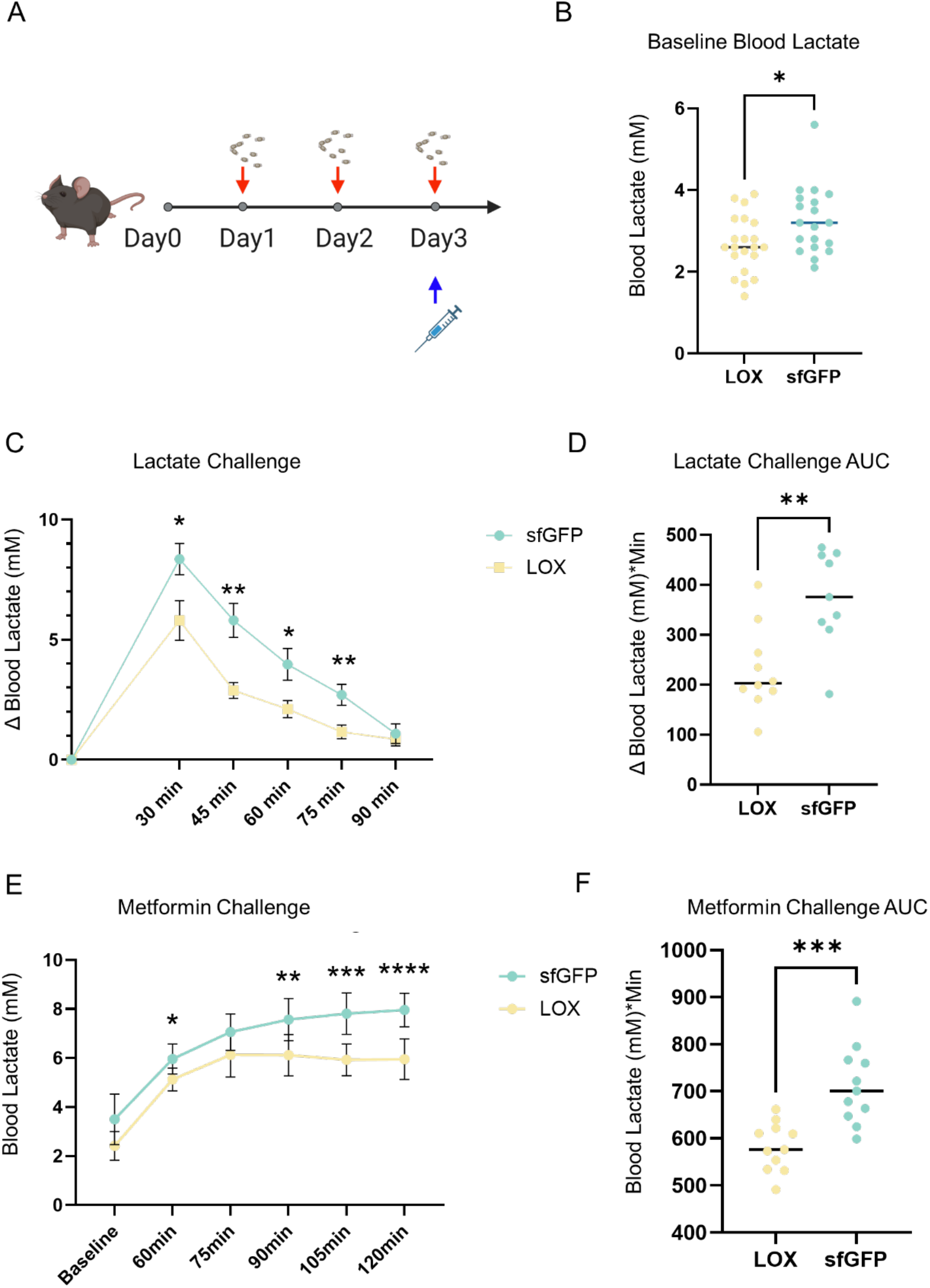
Oral supplementation of *B. subtilis^LOX^* spores reduced baseline blood lactate and accelerated lactate clearance in two mouse models of metabolic challenge. (A) Schematic diagram of the challenge procedures. After daily oral gavage of spores for three days, L-lactate or metformin was injected intraperitoneally on day 3 to increase blood lactate. (B) Combined baseline blood lactate values prior to challenge in both experiments. n= 21 and 19 for *B. subtilis^LOX^* and *B. subtilis^sfGFP^*, respectively. (C) Time course measurements of blood lactate concentration in response to injection of L-lactate. (D) Area Under the Curve (AUC) for blood lactate concentration after injection. n=10 and 9 mice for *B. subtilis^LOX^* and *B. subtilis^sfGFP^*, respectively. (E) Time course measurements of blood lactate concentration in response to metformin challenge. (F) AUC for metformin challenge. n=11 and 10 for *B. subtilis^LOX^* and *B. subtilis^sfGFP^*, respectively. * *p*<0.05, ** *p*<0.01, *** *p*<0.001, **** *p*<0.0001. When not plotted as individual values, data are mean ± SEM. Data were analyzed by student’s T test and repeated measures ANOVA followed by pairwise comparisons with the Sidak correction. Δ Blood Lactate=current lactate concentration-baseline concentration.

In addition to the exogenous administration of sodium lactate in mice, we further assessed *B. subtilis*^LOX^ in a mouse model of drug-induced acute mitochondrial dysfunction using metformin. Metformin is a mitochondrial complex I inhibitor that is known to upregulate serum lactate in humans and mice (35). Utilizing the same gavage schedule as the lactate challenge (**Figure 3A**), mice were administered with an intraperitoneal injection of 400 mg/kg of metformin after the final dose of spores. Blood lactate was measured every 15 minutes between one- and two-hour post-injection. Consistent with the lactate challenge, we observed lower blood lactate values in *B. subtilis*^LOX^-treated mice at 60-, 90-, 105-, and 120-min post-injection (**Figure 3E**), along with a significantly reduced total AUC (*p*=0.01, **Figure 3F**) in comparison to the *B. subtilis*^sfGFP^ group. Collectively, these findings indicate that oral administration of *B. subtilis*^LOX^ spores can promote resilience to lactic acidosis and/or metabolic challenge in both exogenous lactate challenge and pharmacologic inhibition models.

### Oral administration of *B. subtilis* PY79 did not disrupt gut microbiome composition, liver function, or immune homeostasis

To understand whether feeding mice with engineered *B. subtilis* strains caused systemic toxicity, blood samples were collected at 0 hour (before oral gavage), 24 hours, 72 hours, and Day 7 after 1^st^ oral gavage for different biochemical assays (**Figure 4A**). We found that aminotransferase (ALT) and aspartate aminotransferase (AST) levels in both groups were not significantly different from those in the PBS group (**Figure 4B**). One mouse in the LOX group exhibited an elevated AST level. However, it is important to note that the observed levels for both ALT and AST are considerably lower than the upper limits of the reference ranges for healthy mouse serum, which are 24.3 - 115.25 U/L for ALT and 39.55 - 386.05 U/L for AST (35). In parallel, inflammatory cytokine and chemokine levels were quantified by the Luminex Assay. Compared to the PBS group, administration of *B. subtilis*^LOX^ or *B. subtilis*^sfGFP^ did not elevate representative pro-inflammatory factors including IL-1β, monocyte chemoattractant protein 1 (MCP1) and IFNγ (**Figure 4C**). Other anti-inflammatory and pro-inflammatory cytokine levels were also similar to those in the PBS group (**Figure S4**). Moreover, the concentration of TNFα was extremely low and was undetected in all samples. These findings strongly suggest that feeding *B. subtilis* did not induce any discernible side effects in mice. Subsequently, we assessed whether feeding *B. subtilis* would influence the overall composition of the gut microbiome (**Figure 4D**). *B. subtilis* was detected in feces one day after the initial oral gavage and increased with subsequent administrations. However, the total amount of *B. subtilis* reduced 7 days after the first gavage, likely attributed to the cessation of *B. subtilis* administration and the lack of ability to colonize the mouse gut by probiotics in general. By 15 days after the initial gavage, *B. subtilis* was no longer detectable in fecal samples, indicating that *B. subtilis* does not establish long-term colonization in the gut in C57BL/6 mice (**Figure 4E**). Furthermore, there were no discernible differences in colonization between the LOX and sfGFP groups, indicating that the expression of LOX does not likely impact the fitness of *B. subtilis* in the gut (**Figure 4E**). The same stool samples were also collected at the designated time points for 16S rRNA analysis. In terms of relative abundance, *Bacillus* was absent in the top 10 most abundant genera (**Figure S5**). Measures of α-diversity such as Chao1 (**Figure 4F**) or Shannon indices (**Figure S6**) revealed no influence of treatment administered (*p*=0.488 and 0.16 for Shannon and Chao1, respectively) or time point of sample collection relative to gavage (*p*=0.137 and 0.67 for Shannon and Chao1, respectively). Additionally, pairwise adonis testing of weighted UniFrac distances between treatment groups across different time points (Day 1, 3, and Day 7) after the first gavage revealed no difference in β-diversity between the LOX, sfGFP and PBS groups (p>0.05) (**Figure 4G**). This indicates that even at relatively low abundance, engineered *B. subtilis* can introduce the functional impact of LOX on blood lactate levels without significantly altering the microbial community in the host.

**Figure 4.**
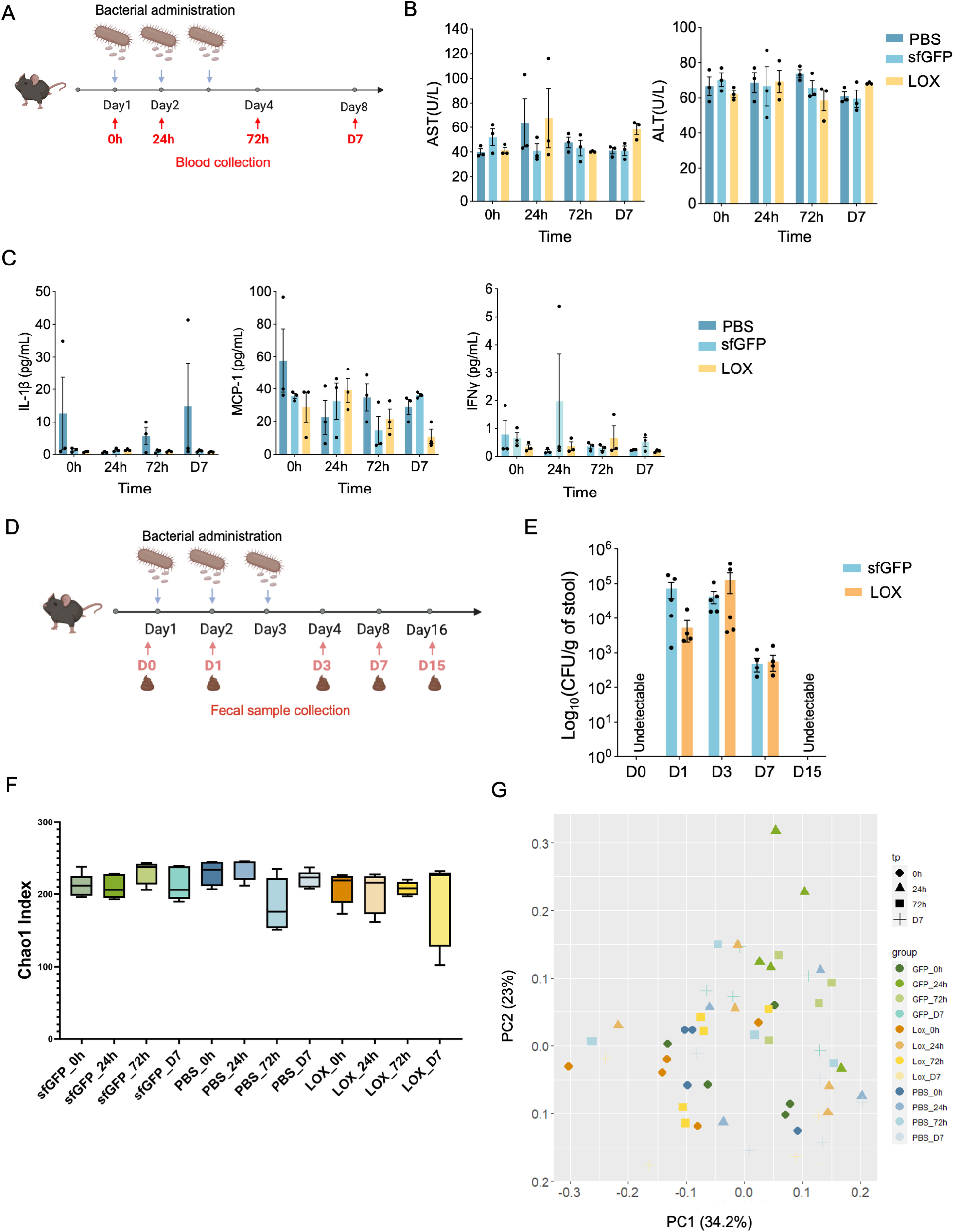
Oral administration of engineered *B. subtilis* PY79 did not disrupt liver function, immune homeostasis, or gut microbiome composition. (A) Schematic of time points for blood collection. “0h” refers to blood samples collected right before the first oral gavage to serve as the baseline. (B) Liver function is represented by circulating ALT and AST levels. ALT and AST concentrations in the plasma of healthy mice range from 24.3 - 115.25 U/L and 39.55 - 386.05 U/L respectively. n=3 mice for each treatment. Data are means ± SEM. (C) Concentrations of proinflammatory cytokines and chemokines in mouse plasma. IL, interleukin; MCP, monocyte chemoattractant protein; IFN, interferon. n=3 mice for each treatment. Data are means ± SEM. (D) Schematic of the experimental procedure for oral gavage and fecal sample collection. “D0” refers to fecal samples collected right before the first oral gavage to serve as the baseline. (E) Colony forming unit (CFU) of *B. subtilis* per gram of feces in mice at respective time points. n=5 mice per treatment per time point. (F) Chao1 index of Alpha diversity for respective treatments and time points. n=5 mice for each treatment. (G) Weighted Unifrac Principal Coordinate Analysis (PCoA) found no differences in community composition between different time points or treatment groups. n=5 mice for each treatment, *p* > 0.05.

## Discussion

The present work establishes a convenient, safe, and effective method to promote accelerated bodily lactate clearance to manage lactic acidosis. We first determined the optimal design *in vitro,* following which we demonstrated that oral administration of the resultant PY79 strain expressing lactate oxidase led to a safe and transient residence in the fecal microbiota. Functionally, this strain accelerated lactate clearance *in vivo* in experimental models of acute lactic acidosis. Not only is lactic acidosis associated with various chronic conditions such as mitochondrial dysfunction, neurodegeneration, liver disorders and diabetes, it is also considered an indicator for clinical mortalities such as hypoxia, cardiac failure, and sepsis (37–39). Given the reliance on concentration-dependent regulation of the LDH reaction, the removal of lactate and the addition of pyruvate has been proven to alleviate tissue-level reductive stress and promote the assimilation of electrons (13). A technology that can accomplish this through simple oral administration may prove beneficial in acute clinical care, as it circumvents many obstacles associated with environments or dosing needed for other currently approved therapies.

Chronically speaking, exposure to elevated lactate concentrations over long periods is associated with the development of various diseases and pathologies. At the cellular level, chronic exposure to elevated lactate has been shown to induce reductive stress, metabolic inflexibility, cellular senescence, and cancer malignancy (40–43). Since elevated blood lactate is purported to represent the mitochondrial status and metabolic flexibility of the respective individual, as previously stated, the clearance of bodily lactate and provision of additional pyruvate may serve to counteract this effect by regulating redox homeostasis through the LDH reaction (13). One limitation of the present work is that the lactate challenges used were merely experimental models with limited external validity. It will be necessary to test more practically relevant metabolic challenge conditions such as treadmill exercise or other models of mitochondrial disease to assess its functional efficacy. Second, blood lactate is the only biochemical outcome measured in this work. It will be important to characterize the effects on not only the metabolome in blood and tissues but also the metabolic and redox status of relevant tissues to unequivocally determine that treatment is conveying a beneficial effect on recipient health and may have a beneficial effect on disease states associated with mitochondrial dysfunction. Given that mitochondrial dysfunction is observed in nearly every significant source of chronic disease-associated mortality, this line of work represents a highly impactful opportunity to help limit healthcare and disease burden from a plethora of sources.

In the current work, we chose *B. subtilis* as our chassis not only due to the practical advantages associated with the resistance to harsh gastric conditions and long shelf life of *B. subtilis* spores, but also because oral administration of *B. subtilis* has been demonstrated to have positive impacts on human gut health. For example, it enhances the function of the host immune system, provides protection against infections, and produces nutrients with high bioavailability (44). Additionally, it has previously been used in commercial products, such as Zbiotic, which is a probiotic designed to remove the toxic byproducts of alcohol metabolism (45). Our proof-of-concept work opens possibilities for engineering other bacterial chassis for reducing lactate accumulation, presenting an alternative strategy in sports medicine, metabolic disorders, and gut health. Finally, while we focus on engineering *B. subtilis*, it could provide added benefit to employ other existing bacterial chassis for this purpose such as *E. coli Nissle* and genetically tractable commensal bacteria which can colonize over long periods. Models with transient or long-term colonization present separate advantages from a treatment standpoint and may complement the current model nicely. While it appears that the consumption of extracellular lactate does not require the overexpression of a lactate importer, it remains to be addressed whether the co-expression of a lactate importer may further improve the efficiency of lactate catabolism by facilitating enhanced uptake.

In conclusion, we have demonstrated that *B. subtilis* PY79 is a feasible, transient, and safe delivery vessel for therapeutic enzymes, such as lactate oxidase, that achieve targeted modulation of systemic metabolome through its short-term residence in the gut. Additionally, given its ability to form spores that are resilient to harsh conditions and can germinate in the gut, this represents a translatable platform that can be easily stored, delivered, and used for a variety of dosing schedules and pathologies. Due to prior research connecting extracellular lactate on intracellular redox and clinical outcomes, the observed decreases in lactate appear to possess clinically significant potential but warrant further investigation into functional and intracellular outcomes to test this hypothesis.

## Materials and Methods

### Chemicals and Reagents

All chemicals and bacterial culture broth were purchased from Fisher Scientific International Inc. (Cambridge, MA, USA) unless otherwise noted, and were of the highest purity or analytical grade commercially available. All molecular cloning reagents including restriction enzymes, competent cells, and the HIFI assembly kit were purchased from New England Biolabs (Ipswich, MA, USA). DNA oligonucleotides were ordered from Sigma Aldrich (St. Louis, MO, USA). LOX, CAT, and UtCAT gblocks were ordered from IDT with *B. subtilis* codon optimization.

### B. subtilis General Growth Conditions

*B. subtilis* strains were grown at 30°C, 220rpm overnight in LB medium with or without 5ug/mL chloramphenicol. Overnight bacterial culture was diluted 10 times followed by 37°C 220rpm culturing until the OD_600_ reached between 0.6 and 0.8. For *in vivo* studies, bacteria were stored in formulation buffer (2.28g/L KH_2_PO_4_, 14.5g/L K_2_HPO_4_, 15% glycerol, pH 7.5) at 10^11^ CFU/mL at -80 °C.

### Constitutive Strain Construction

All *B. subtilis* strains were constructed in WB800N first. Primers can be found in Supplementary Table 1. All constructs were confirmed by Sanger sequencing.

All *amyE* locus integration fragments contain three parts: *amyE* 5’ arm with *P_NBP3510_*, GOI, and *amyE* 3’ arm with chloramphenicol resistance. The integration fragment was assembled from different plasmids and chromosomal DNA of *B. subtilis* to make *ΔamyE::cat*_sfGFP first which was used as a new template to later make LOX constructs. FLAG and HA tag were added at 3’ of LOX and UtCAT respectively. For fusion enzyme CAT_L20_LOX and UtCAT_L20_LOX, FLAG and HA were added at 5’ of CAT and 3’ of UtCAT respectively.

*ΔamyE::cat*_sfGFP: The *amyE* 5’ arm was amplified through WB800N genomic DNA with the primers B685_V2/B742. *P_NBP3510_*_sfGFP was amplified from WBSGFP with B741/B747. AmyE 3’ arm was amplified through lab plasmid pHT01-NBP3510-sfGFP with B746/B690. Three fragments were first combined through HIFI assembly kit followed by cementing PCR via primers B685_V2/B690 before transformation into *B. subtilis* as described below.

*ΔamyE::cat* _LOX: *amyE* 5’ arm with *P_NBP3510_* and with *amyE* 3’ arm with Cm^r^ were amplified by B685_V2/B774 and B776/B690 separately through the chromosomal DNA of PY79 *ΔamyE::cat*_sfGFP. LOX was amplified via B775/B777.

*ΔamyE::cat*_CAT_L20_LOX: *amyE* 5’ arm with *P_NBP3510_* and *amyE* 3’ arm with Cm^r^ were amplified by B685_V2/B781 and B776/B690 separately through chromosomal DNA of PY79 *ΔamyE::cat*_sfGFP. CAT was amplified via B782/B218 through Gblock. LOX was amplified via B218/B783

*ΔamyE::cat*_LOX_B0035_UtCAT: *amyE* 5’ arm with LOX was amplified via B685_V2/B614 through chromosomal DNA of PY79 *ΔamyE::cat*_LOX. B0035_UtCAT was amplified via B617/B780 through UtCAT Gblock. *amyE* 3’ arms with Cm^r^ were amplified by B776/B690.

*ΔamyE::cat*_LOX_R0 _UtCAT: *amyE* 5’ arm with LOX was amplified via B685_V2/B778 through chromosomal DNA of PY79 Δ*amyE*::cat_LOX. R0_UtCAT was amplified via B779/B780 through UtCAT Gblock. *amyE* 3’ arms with Cm^r^ were amplified by B776/B690.

*ΔamyE::cat*_UtCAT_L20 _LOX: *amyE* 5’ arm with *P_NBP3510_* and *amyE* 3’ arm with Cm^r^ were amplified by B685_V2/B784 and B776/B690 separately through chromosomal DNA of PY79 *ΔamyE::cat*_sfGFP. UtCAT was amplified through gblock via B785/B786. L20_LOX was amplified through *ΔamyE::cat*_CAT_L20_LOX via B787/B783.

### B. subtilis transformation

Cementing PCR products were cleaned up by PB buffer before transformation. MC media was made at a 10x stock with final concentration of 14.036% K_2_HPO_4_ (w/v), 5.239% KH_2_PO_4_ (w/v), 20% glucose (w/v), 30 mM trisodium citrate, 0.022% Ferric ammonium citrate (w/v), 2% potassium glutamate (w/v), 1% casein digest (Difco). The 10xMC stock was filtered and stored at -20 °C. To make1xMC media, supplemented 1xMC with 3 mM MgSO_4_. *B. subtilis* strains were re-streaked one day before transformation. Single colonies were inoculated into 1 ml 1xMC media followed by 37 °C, 220rpm shaking for 3-4h until turbid. Cementing PCR products were purified by PB buffer and directly added into 300ul of culture. Culture shaking was resumed at 37 °C for another 2.5 h followed by centrifugation to get rid of 250 μL supernatant. The remaining culture was plated on the LB plates with 5 μg/mL Chloramphenicol. Multiple single colonies were picked the next day and restreaked on the selective LB plates to ensure integration. Colonies which grew successfully were picked and verified by Sanger sequencing.

### SPP1 Phage Transduction

The donor colony was inoculated in 3 mL TY broth (10g tryptone, 5g yeast extract, 5g NaCl per liter) and grew at 37 °C until the culture was turbid. SPPI phage dilutions were prepared from 10^-2^ to 10^-6^ in sterile tubes. 100 μL of each dilution was added into a clean tube, followed by a 200 uL turbid donor culture. Mixtures were incubated statically at 37 °C for 15 mins. TY soft agar (5g agar/L) was melted during the incubation and supplemented with 10 mM MgSO_4_ and 0.1 mM MnSO_4_ as working concentration. 3 mL soft agar was added to the bacterial and phage mixture, and the mixture was vortexed. Tube contents were dumped onto the TY agar plate (15g agar/L) and the plate was swirled around to make sure the mixtures covered the entire plate surface. Plates were dried and incubated overnight at 37 °C. The next day lysates were harvested by adding 5 mL TY media to appropriate plaques. Soft agar and TY broth were scraped and incubated at RT for 10 mins, then spun down at 4000 rpm for 15 mins. Lysate was filtered using 0.45 µm syringe filter into a new tube and 100 μL chloroform was added for storage. For transduction experiments, lysates were prepared from the donor strains as described above. The recipient colony was inoculated in 3 mL TY broth followed by shaking at 37 °C and 250 rpm until dense.1ml bacterial culture was mixed with 10 μL phage stock followed by adding 9ml TY broth. The mixture was statically incubated at 37 °C in a water bath for 30 mins. The mixture was then centrifuged at 4000 rpm for 20 mins. Most of the supernatant was discarded, and the pellet was resuspended in the remaining volume. 100ul bacterial culture was plated on selective media with 10mM sodium citrate. Plates were then incubated overnight at 37 °C. The next day, 3-4 different colonies were restreaked onto selective plates without sodium citrate.

### Catalase Activity Measurement

Relative catalase activities between different engineered strains were measured and compared by following a previously published protocol (33). Briefly, 1.6x10^9^ bacteria were pelleted and resuspended in 100 uL 0.9% NaCl and 100 μL 1% Triton X-100. The solution was transferred to FACS tubes. Subsequently, 100 μL 3% hydrogen peroxide solution (Sigma, catalog #88597-100ML-F) was added to the solution followed by thorough mixing. After the completion of each reaction, the height of the oxygen-forming foam was recorded by subtracting the height of the original solution.

### Western Blotting

200 uL bacterial pellet was collected by centrifugation and resuspended in 50 µL 1xTE (10mM Tris-HCl, 1 mM EDTA, pH8.0) with 1.5 μL 50mg/ml lysozyme followed by 10 mins incubation in 37 °C water bath. 10ul 6xSDS loading dye was added to the suspension. The samples were first separated by 12% SDS-PAGE gel and then transferred to a nitrocellulose membrane (Fisher Scientific). The membranes were incubated with the Anti-FLAG epitope (DYKDDDDK, Biolegend, San Diego, CA, catalog# 637301) or Anti-HA epitope (ABclonal, Woburn, MA catalog# AE008) with 1:2000 dilution overnight in a cold room, and the secondary antibody anti-rat IgG HRP (Cell Signaling Technology, Danvers, MA, catalog# 7077) for Anti-FLAG or anti-mouse IgG HRP (Invitrogen, Waltham, MA, catalog# 626520) for Anti-HA with the same dilution at room temperature for 1 h. Premixed Pierce^TM^ 3,3’diaminobenzidine (DAB) substrate (Fisher Scientific) was used to detect the target proteins.

### In Vitro Lactate Depletion Measurement

*Bacillus subtilis* LOX, and sfGFP strains were inoculated in LB with 5 µg/mL Cm at 30 °C overnight. Overnight cultures were diluted 10 times into 15 mL fresh LB with 5 µg/mL Cm followed by incubation at 37 °C until OD_600_=0.8. Pellets were collected by centrifugation at 4000 rpm 20°C for 10 mins and washed with PBS three times. 100 uL 0.5 M perchloric acid (PCA) was pre-aliquoted in a 96-well V bottom plate and placed on ice. The pellet was resuspended with PBS and the OD_600_ was adjusted to 1. Bacterial culture and sodium-L-lactate (Fisher, catalog# AAL1450006) standards at concentrations of 5 mM, 2.5 mM, 1.25 mM, 0.625 mM, and 0 mM were added to the corresponding positions of the layout in a 2 mL deep well plate. Sodium-L-lactate was added to the corresponding bacterial suspension with 5 mM working concentration. 100 uL reaction mixture was mixed and transferred to pre-aliquoted PCA on ice as the baseline. The 96-well deep well plate was kept in a shaker at 37 °C 220rpm to collect samples at desired timepoints. The V-bottom plates were spun down at 4 °C 4000rpm for 10 mins after 30mins incubation on ice. 150 uL supernatant was transferred into a new 96-well cell plate and neutralized by 30 µL 2.3M KHCO_3_ followed by overnight precipitation. The plates were centrifuged at 4000rpm 4 °C the next day for 10 mins to collect the samples for L-lactate detection. For L-lactate detection, 10 uL of each sample and water blanks were added to 96-flat black well plates with three technical repeats. 185 µL LA detection buffer (100 mM Hydrazine dihydrochloride, 100 mM glycine, 0.5 mM NAD^+^, pH 10) was added to each well followed by 15 mins incubation. The plate was read at excitation 340, emission 460 as the baseline. 5 μL 8U/mL L-Lactic Dehydrogenase (LDH) from rabbit muscle Type II, ammonium sulfate suspension (Sigma L2500) was added to all the wells followed by 250rpm shaking for 5 mins to mix well, and plates were kept in the dark. Plates were then incubated in the dark for 2 hours and read.

### Spore Preparation

Bacterial culture was plated on selective 2xSchaeffer’s medium-glucose plates followed by 1-week 37 °C incubation(^46^). Spores were scraped from the plate to centrifuge tubes and were then washed by resuspension in ddH_2_O. Spores were then pelleted via centrifugation at 5000rpm for 10 mins at RT. The pellet was then resuspended in ddH_2_O followed by three cycles of 1 min sonication at 20 W and 2,100 Joules, and 1 min rest. Spores were subjected to 1 min sonication followed by 1 min rest. This cycle was repeated step three times a day for 3 days. Spores were aliquoted at 10^9^ CFU/mL each tube and stored at -80 °C for later use.

### In Vivo Lactate Induction Experiments

Five female C57BL/6 mice (Jackson Laboratories) were housed in a single cage for each condition. 6 to 8-week-old mice were gavaged daily with 100 µL containing 10^11^ CFU/kg body weight engineered PY79 spores of the corresponding strain for 3 days. 20 mins after the 3^rd^ oral gavage, baseline blood lactate level was measured by Lactate Plus: Blood Lactate Measuring Meter-Version 2 (NovaBiomedical) and Lactate Plus Meter Test Strips (NovaBiomedical, catalog# 40813) through tail snipping. Sodium-L-lactate was administered via intraperitoneal injection at a dose of 2.3 g/kg body weight. Blood lactate was measured 30, 45-, 60-, 75-, and 90-min post injection. For the metformin challenge, 20 mins after the final gavage, baseline blood lactate was measured, and mice were intraperitoneally injected with 400 mg/kg metformin (Fisher Scientific, Waltham MA) in 500 µL PBS. Blood lactate measurements were taken at 60, 75, 90, 105, and 120 minutes. Values from both separate experiments are included in the figure containing baseline values (**Figure 3B**).

### In Vivo Colonization and Safety Experiments

All animal experiments were approved by the University of Michigan and Northeastern University Institutional Animal Care and Use Committee (IACUC). Five female C57BL/6 mice (Jackson Laboratories) were housed in a single cage for each condition. 6 to 8-week-old mice were gavaged daily with PBS or 200 µL engineered *B. subtilis* (10^10^ vegetative cells plus 10^8^ spores) for 3 days. Baseline (0h) blood samples were collected into BD Microtainer™ Capillary Blood Collector containing EDTA, right before the 1^st^ oral administration of spores followed by blood collection at 24 h, 72 h, and 7 days post-1^st^ oral gavage. Plasma was isolated by centrifugation at 3000xg for 30 mins at 4 °C. The plasma was transferred to clean 0.65 mL tubes for Luminex Assay (MDF10) through Eve Technology (Canada). Plasma was stored in -80 °C until use. AST and ALT levels were sent to the University of Michigan IVAC facility for mini liver chemistry panel (MI1015) analysis. Stools were collected before 1^st^ gavage, and 1 day, 3 days, 5 days, 7 days, 15 days, 18 days after 1^st^ gavage. Fecal samples were placed in 300 uL sterile PBS buffer followed by 15 mins incubation. The mixture was homogenized by micro-tissue homogenizer. The solution was obtained by centrifugation and applied to LB agar with 7.5ug/mL chloramphenicol, 0.25ug/mL gentamycin, and 0.25 μg/mL erythromycin. Plates were incubated at 37 °C for 12-16 hours. CFU was determined by standard CFU counting method.

### Genomic DNA Extraction from Feces

C57BL/6 mouse feces were collected before 1^st^ gavage, and 1 day, 3 days, 5 days, and 7 days, after 1^st^ gavage. Feces were immediately frozen in liquid nitrogen and saved in -80 °C for later use. Genomic DNA was extracted by Quick-DNA Fecal/Soil Microbe Miniprep Kit (Zymo, catalog# D6010) following the instructions.

### 16S rRNA Gene Sequencing and Analysis

16s rRNA analyses were performed both in-house and by Novogene, China. gDNA concentration was measured with 1% agarose gels. According to the concentration, DNA was diluted to 1ng/μL using sterile water. 16S rRNA V4 specific primers are 515F (5’-GTGCCAGCMGCCGCGGTAA-3’) and 806A (5’-GGACTACHVGGGTWTCTAAT-3’). All PCR reactions were carried out in 30 μL reactions with 15 μL of Phusion® High-Fidelity PCR Master Mix (New England Biolabs), 0.2 μM of forward and reverse primers, and about 10 ng template DNA. Thermal cycling is started with the initial denaturation at 98°C for 1 min, followed by 30 cycles of denaturation at 98°C for 10 s, annealing at 50°C for 30 s, and elongation at 72°C for 60 s. Finally, within 72°C for 5 min. PCR products were mixed with an equal volume of SYB Green loading buffer and operated electrophoresis on 2% agarose gel for quantification and qualification followed by purification with GeneJET Gel Extraction Kit (Thermo Scientific). Sequencing libraries were generated using NEB Next® Ultra^TM^ DNA Library Prep Kit for Illumina (NEB, USA) following the manufacturer’s recommendations and index codes were added. The library quality was assessed on the Qubit@2.0 Fluorometer (Thermo Scientific) and Agilent Bioanalyzer 2100 system. At last, the library was sequenced on an Illumina HiSeq 2500 platform, generating 250 bp paired-end reads. For taxa relative abundance outcomes, qualified reads were filtered by QIIME software to obtain clean tags. All tags were compared with the reference database using UCHIME algorithm to obtain effective tags only which were used for diversity analysis. For diversity analyses, FASTA files were processed with DADA2 for R, with primer sorted and demultiplexed sequences filtered and trimmed at truncation lengths of 240 and 160 bp for forward and reverse reads respectively with a minimum quality score of 20. Following merging and removing of chimeric reads, taxonomy was assigned as Amplicon Sequence Variants (ASVs) with Silva v138 taxonomic database. Using phyloseq for R, ASVs not present at least 5 times in 50% of samples were pruned, and in all outcomes aside from alpha diversity, reads were rarefied to even sampling depth.

### Statistical Analysis

All statistical analyses were performed using GraphPad Prism 9 (San Diego, CA, USA). Data were analyzed with one-way or two-way analysis of variance (ANOVA) for statistical significance.

## Acknowledgments

This work was supported by NIBIB (R21EB030769), CDMRP PRMRP (W81XWH-21-1-0016), and in part by NIDCR (R01DE030943). WBSGFP was kindly provided by Prof. Xin Yan from Nanjing Agricultural University, China. SPP1 phage was a generous gift by Professor Dan Kearns from Indiana University Bloomington. We would like to express our gratitude to the ULAM Pathology Core at the University of Michigan for liver panel analyses; Professor Sara Rouhanifard at Northeastern University’s Department of Bioengineering for generously sharing her lab’s plate reader with us; Professor Ke Zhang at Northeastern University’s Department of Chemistry for sharing his lab’s equipment and space. We also wish to thank my colleague Morgen Benson for her contribution to the sfGFP construct during the lab rotation.

## Author contributions

JL conceived the ideas and supervised the experiments. MY designed and conducted the experiments. NH assisted with 16S rRNA α- and β-diversity analysis and conducted experiments validating *in vivo* efficacy. NY assisted with the animal work. JY assisted with material preparation and plate reading for LOX activity *in vitro* tests, as well as phage transduction. MG assisted with spore preparation and western blotting. ZW prepared the *B. subtilis* vegetative cells and assisted in measuring blood lactate. PC provided guidance for mice restraint and blood collection. SY provided guidance for mice oral administration. JC developed the *in vitro* lactate assay. All authors contributed to the discussion of the results and the writing and editing of the manuscript.

