## Supplemental Figures for "Engineered *Bacillus subtilis* as oral probiotics to enhance clearance of blood lactate"

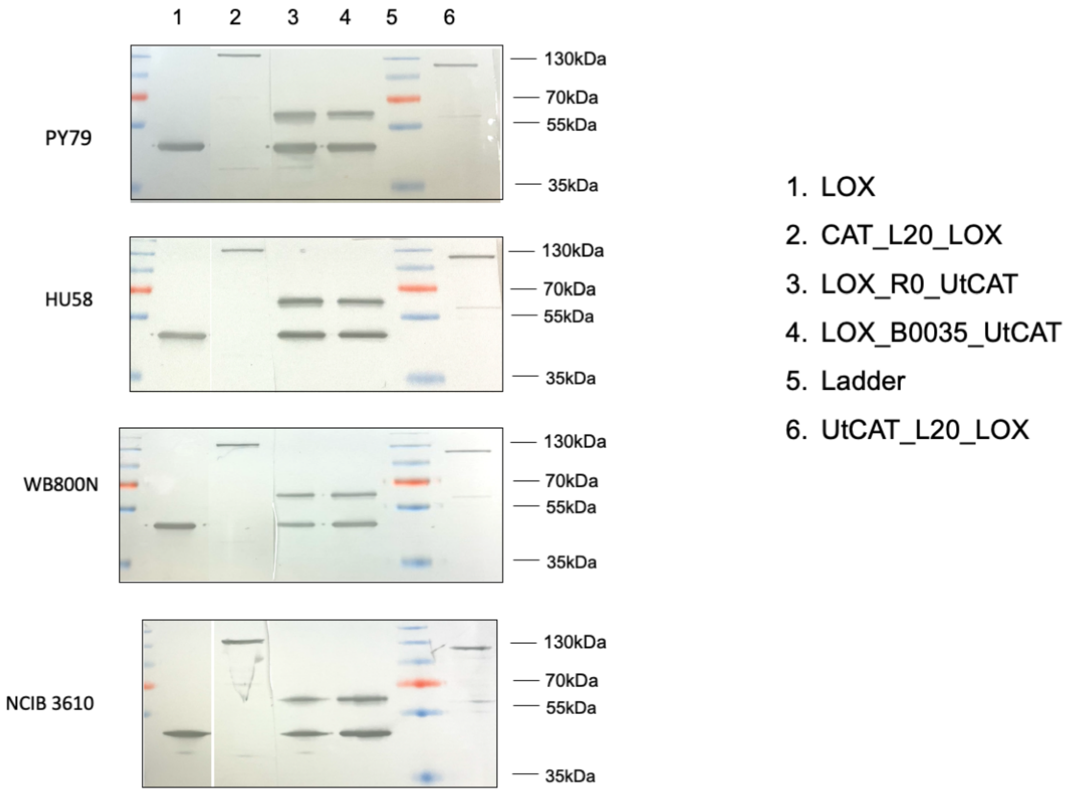

**Supplementary Figure 1: Expression of recombinant enzymes in various *B. subtilis* strains**. The predicted molecular weights for LOX, UtCAT, CAT_L20_LOX (fusion protein), UtCAT_L20_LOX (fusion protein) are 41kDa, 59kDa, 120kDa, and 101kDa. The data are representative of at least 3 biological repeats with similar results. The membranes were cut and merged because some Western blotting experiments were performed on different days.

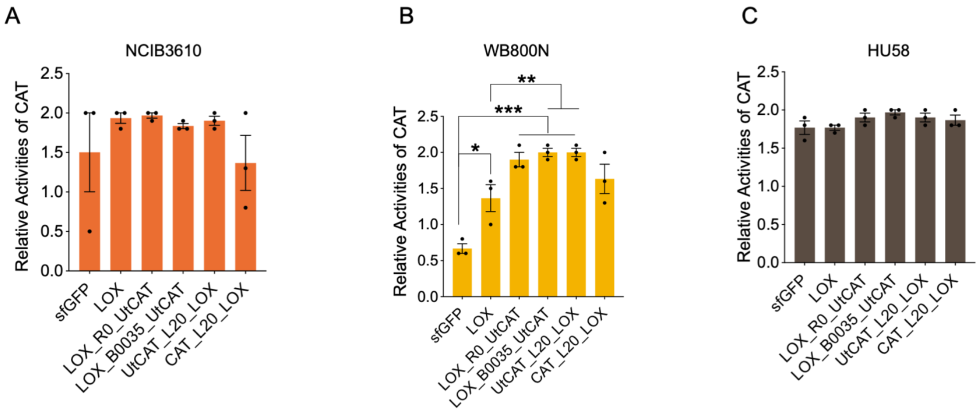

**Supplementary Figure 2: The catalase activities of *B. subtilis* strains (A) NCIB3610, (B) WB800N and (C) HU58 encoding different constructs.** The breakdown of hydrogen peroxide into oxygen was catalyzed by endogenous catalase (“sfGFP” or “LOX”) or in combination with *E. coli* catalase (CAT) or *U. thermosphaericus* catalase (UtCAT) for “LOX_R0_UtCAT”, “LOX_B0035_UtCAT”, “UtCAT_L20_LOX” and “CAT_L20_LOX”. N=3 independent experiments. Data were analyzed by unpaired t-test, *$p < 0.05$, **$p < 0.01$, ***$p < 0.001$. Data are mean ± SEM.

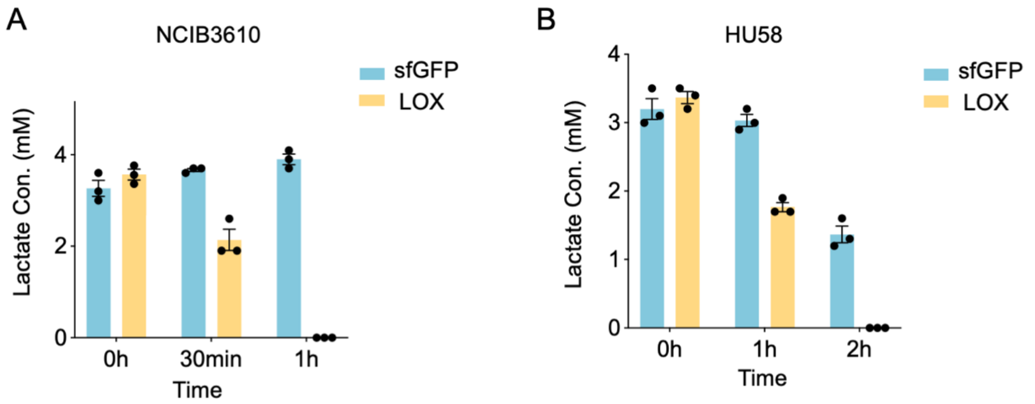

**Supplementary Figure 3**: (A) NCIB3610 and (B) HU58 *B. subtilis*$^{LOX}$ consumed exogenous L-lactate within 1 and 2 hours, respectively, compared to *B. subtilis*$^{sfGFP}$. N=3 independent experiments and n=3 technical replicates. Data are mean ± SEM.

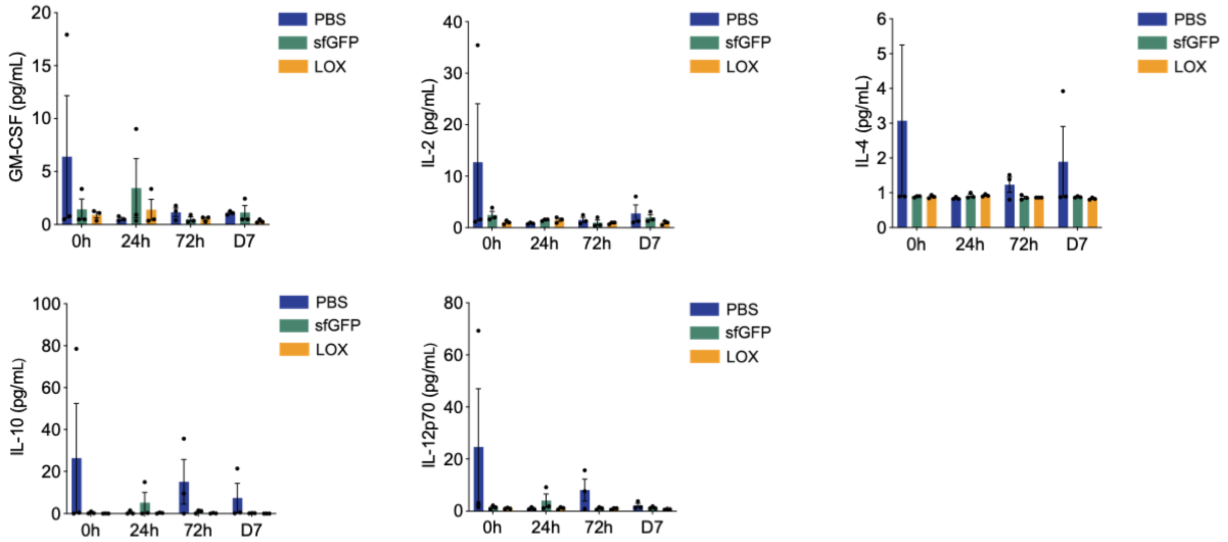

**Supplementary Figure 4:** Profiling of cytokines (GM-CSF, IL-2, IL-4, IL-10 and IL-12p70) in the plasma of treated mice indicates a lack of side effects. Cytokines were quantified by the Luminex Assay. “0h” refers to blood samples collected before the first oral gavage to serve as the baseline. n=3 mice for each treatment group.

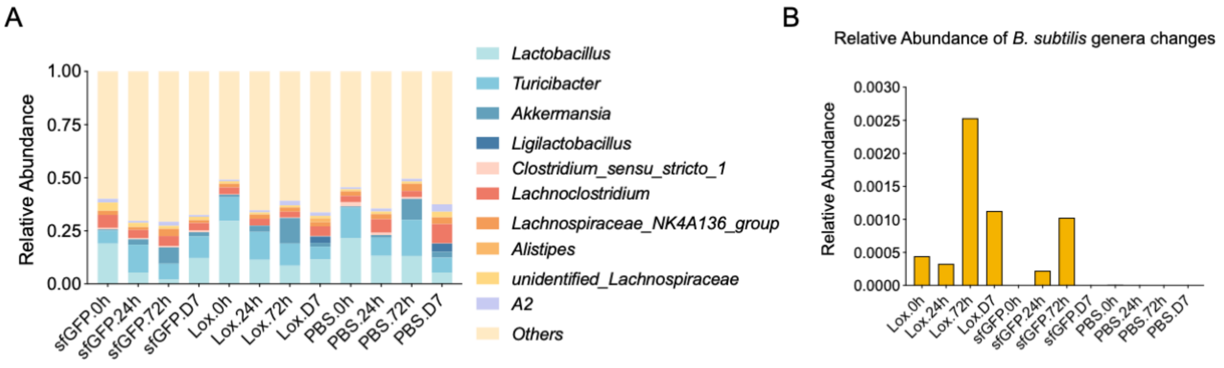

**Supplementary Figure 5:** (A) Relative abundance of the top 10 genera in each treatment group across time points indicates that *Bacillus* is not among 10 most abundant genera. (B) Relative abundance of *Bacillus* based on the 16S sequencing. “0h” refers to when fecal samples were collected right before the first oral gavage to serve as the baseline. n=5 mice for each treatment.

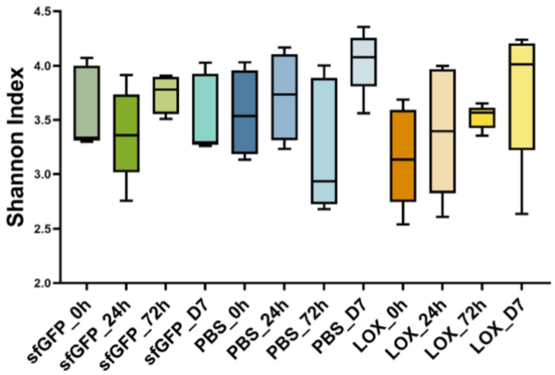

**Supplementary Figure 6:** Shannon index of α-diversity analysis suggests no significant difference in community evenness between *B. subtilis* and PBS treatment groups in C57BL/6 mice. $p$=0.488 and 0.137 for influence of treatment and time point, respectively. n=5 mice for each treatment.

**Supplemental Table 1: primers used in this study**

| Primer | Sequences 5’ to 3’ | Description | Construct |
|---|---|---|---|
| B685_V2 | TGCTTACGATGTACGACAGGGGG | amyE_up_fwd | *ΔamyE ::cat*_sfGFP |
| B742 | CTAAAAATTTGTAATTAAGAAGGAGTGATTACcgatcagaccagtttttaatttg | amyE_up_rev | |
| B741 | GTAATCACTCCTTCTTAATTACAAATTTTTAG | $P_{NBP3510}$_sfGFP_fwd | |
| B747 | GCGTAAGGAAATCCATTATGTACTATTTttcatatatttggctgcggtag | $P_{NBP3510}$_sfGFP_rev | |
| B746 | AAATAGTACATAATGGATTTCCTTACGC | *cat*_amyE_down_fwd | |
| B690 | CATCCTTGCAGGGTATGTTTCTCTTTG | amyE_cat__down_rev | |
| B774 | tcaatgtcattgttattcatACGTTCTACCTTTGTCAAAC | amyE_$P_{NBP3510}$_rev | *ΔamyE ::cat*_LOX |
| B775 | AACGTatgaataacaatgacattga | LOX_fwd | |
| B777 | GGACGTCGACTCTAGAttactatttgtcatcgtcg | LOX_rev | |
| B776 | tagtaaTCTAGAGTCGACGTCC | *cat*_amyE_down_fwd | |
| B781 | tcgtcgtctttgtagtccatACGTTCTACCTTTGTCAAAC | amyE_$P_{NBP3510}$_rev | *ΔamyE ::cat*_CAT_L20_LOX |
| B782 | GTatggactacaaagacgacgat | CAT_fwd | |
| B218 | caccttccgagccaccgccctcggagcctcccgcaccacttgcggcaggaattttgtcaatc | CAT_rev | |
| B219 | ggtggctcggaaggtgggacgagcggcgccaccaataacaatgacattgaatataatgc | L20_LOX_fwd | |
| B783 | GGACGTCGACTCTAGAttactagtattcataaccgtatg | L20_LOX_rev | |
| B614 | agtattctcctctttaatctctagattactatttgtcatcgtcg | LOX _rev | *ΔamyE ::cat*_LOX_B0035_UtCAT |
| B617 | agagattaaagaggagaatactagatgaccaatattaatgataaacg | B0035_UtCAT _fwd | |
| B780 | GGACGTCGACTCTAGAttactatgcatagtctggc | B0035_UtCAT _rev | |
| B778 | gtttgtcctccttattagttaatcttactatttgtcatcgtcg | LOX_Rev | *ΔamyE ::cat*_LOX_R0 _UtCAT |
| B779 | gattaactaataaggaggacaaacatgaccaatattaatgataaacg | R0_UtCAT_Fwd | |
| B780 | GGACGTCGACTCTAGAttactatgcatagtctggc | R0_UtCAT_rev | |
| B784 | TTATCATTAATATTGGTCATACGTTCTACCTTTGTCAAAC | amyE_$P_{NBP3510}$_rev | *ΔamyE ::cat*_UtCAT_L20 _LOX |

| | | |
|---|---|---|
| B785 | TAGAACGTATGACCAATATTAATGATAA | UtCAT _fwd |
| B786 | ctcccgcaccacttgcTGCATAGTCTGGCACGTC | UtCAT _rev |
| B787 | TATGCAgcaagtggtgcgggag | L20_LOX_fwd |
